# Coupled small molecules target RNA interference and JAK/STAT signaling to reduce Zika virus infection in *Aedes aegypti*

**DOI:** 10.1101/2021.08.23.457417

**Authors:** Chasity E. Trammell, Gabriela Ramirez, Irma Sanchez-Vargas, Shirley Luckhart, Rushika Perera, Alan G. Goodman

**Author notes:** Correspondence (R.P.); (A.G.G.).

## Abstract

The recent global Zika epidemics have revealed the significant threat that mosquito-borne viruses pose. There are currently no effective vaccines or prophylactics to prevent Zika virus (ZIKV) infection. Limiting exposure to infected mosquitoes is best way to reduce disease incidence. Recent studies have focused on targeting mosquito reproduction and immune responses to reduce transmission. In particular, previous work evaluated the effect of insulin signaling on antiviral JAK/STAT and RNAi in vector mosquitoes. In this work, we demonstrate that targeting insulin signaling through the repurposing of small molecule drugs results in the activation of both of these antiviral pathways. Activation of this coordinated response additively reduced ZIKV levels in *Aedes aegypti* mosquitoes. This effect included a quantitatively greater reduction in salivary gland ZIKV levels relative to single pathway activation, indicating the potential for field delivery of these small molecules to substantially reduce virus transmission.

## INTRODUCTION

Mosquito-borne viruses pose a significant global health threat and this threat is increased by dynamic ecological and human factors. Global warming and urbanization have permitted mosquitoes and arboviruses to spread into previously virus-free regions (Alaniz et al., 2018; Samy et al., 2016). This occurred during the 2015-17 Zika virus (ZIKV) Western hemisphere epidemic that originated in South America and spread into North America, resulting in 538,451 suspected cases, 223,477 confirmed cases, and 3,720 congenital syndrome cases (Pan American Health Organization, 2015, 2016). Subsequent outbreaks have followed that establishes ZIKV as an active pathogen of concern that requires intervention (Bhargavi and Moa, 2020). Current efforts have focused on strategies to reduce virus transmission to and from the mosquito vector, including the use of insecticides and biological/genetic manipulation of primary vector species. The introduction of the bacterial symbiont *Wolbachia* to reduce flavivirus infection in the major arbovirus vector species *Aedes aegypti* (Aliota et al., 2016; Dutra et al., 2016; Haqshenas et al., 2019) and the release of genetically modified individuals to reduce transmission-competent progeny of this species (Waltz, 2021) have been included among the latter strategies. There is evidence to suggest that *Wolbachia*, while effective in reducing ZIKV and dengue virus (DENV) infection in targeted species may inadvertently enhance replication of West Nile virus (WNV) (Dodson et al., 2014). It is also not known how effective or advantageous genetically modified mosquito populations are compared to wild type populations or to other various viruses (Evans et al., 2019; Resnik, 2017). Because of these challenges, additional strategies to reduce vector transmission of these important viral pathogens are critically needed.

As an alternative strategy, it may be possible to reduce or block arborvirus transmission through mosquito-targeted delivery of bioactive small molecules at attractive sugar bait stations, a modification of the successful delivery of toxic baits for mosquito control (Dong and Dimopoulos, 2021). To this end, it is necessary to identify druggable mosquito antiviral effectors and their upstream regulatory factors. The insulin/insulin-like growth factor signaling (IIS) cascade regulates RNA interference (RNAi) and JAK/STAT antiviral immunity against West Nile virus (WNV), dengue virus (DENV), and ZIKV (Ahlers et al., 2019; Trammell and Goodman, 2019). In *Drosophila melanogaster*, the IIS-dependent transcription factor forkhead box O (FOXO) induces expression of RNAi transcripts *AGO2* and *Dicer-2* (Spellberg and Marr, 2015). We demonstrated that ingestion of exogenous insulin reduced expression of these RNAi components in WNV-infected *Culex quinquefasciatus* (Ahlers et al., 2019) and that manipulation of IIS-dependent extracellular-signal regulated kinases (ERK) activation reduced WNV infection in this mosquito host (Ahlers et al., 2019). Further, insulin treatment suppressed the activation of RNAi, while activating ERK-dependent JAK/STAT induction of unpaired (upd) ligands to control WNV replication *in vitro* and *in vivo* (Ahlers et al., 2019). Previous studies established that both JAK/STAT and RNAi antiviral pathways are independently involved in arthropod antiviral immunity to ZIKV (Harsh et al., 2018, 2020; Xu et al., 2019). To date, however, no mechanism(s) have been established whereby both antiviral pathways are induced simultaneously in response to arthropod infection.

In this study, we repurposed small molecules that target IIS pathway proteins to induce simultaneous activation of RNAi and JAK/STAT signaling in *Ae. aegypti*. Specifically, we used the potent insulin mimetic demethylasterriquinone B1 (DMAQ-B1), an activating ligand of the insulin receptor (InR) (Zhang et al., 1999) and Protein kinase B (AKT) inhibitor VIII, which reduces AKT phosphorylation (Lindsley et al., 2005). Small molecule treatment induced activation of JAK/STAT via ERK and blocked inhibition of RNAi via the AKT/FOXO signaling axis. Combined treatment with DMAQ-B1 and AKT inhibitor VIII significantly lowered ZIKV titers in *Ae. aegypti* cells and adult female mosquitoes relative to single treatment and vehicle control. Combined treatment also additively reduced salivary gland virus titers, a surrogate measure of reduced transmission efficacy (Ferguson et al., 2015; Raquin and Lambrechts, 2017). Accordingly, we argue that activation of both antiviral pathways resulted in enhanced defenses that lowered viral titers to non-detectable levels. This work demonstrates the feasibility of strategically targeting mosquito immunity via IIS is a means of reducing a clinically relevant strain of ZIKV infection and transmission at the vector level.

## RESULTS

### *DMAQ-B1 and AKT inhibitor VIII activated* Aedes aegypti *insulin and antiviral signaling pathways*

Since the JAK/STAT and RNAi antiviral pathways are linked to IIS, we sought to test the activity of small molecules against phosphorylation of key IIS protein targets and activity of these antiviral pathways. Given that phosphorylation of AKT and ERK correlate with activation of IIS and JAK/STAT signaling (Ahlers et al., 2019; Boulton et al., 1991), respectively, and that FOXO phosphorylation is consistent with suppression of RNAi (Biggs et al., 1999; Brunet et al., 1999; Spellberg and Marr, 2015), we used these readouts to evaluate the efficacy of DMAQ-B1 and AKT inhibitor VIII control of RNAi and JAK/STAT.

Protein lysates from *Ae. aegypti* Aag2 cells treated with 1% DMSO vehicle, 1 μM DMAQ-B1, 10 μM AKT inhibitor VIII, or these combined drug treatment for 24 hours were analyzed by western blot for phosphorylation of AKT, FOXO, and ERK (**Fig. 1A**). Drug concentrations were based on prior toxicity analysis (**Fig. S1**). DMAQ-B1 treatment was associated with the highest levels of AKT and FOXO phosphorylation; this phosphorylation was significantly reduced when combined with AKT inhibitor VIII (**Fig. 1B-C**). Single drug and combined drug treatments were associated with increased ERK phosphorylation relative to vehicle control (**Fig. 1D**). We validated these findings by immunofluorescence microscopy of 24 h treated cells probed for phospho-FOXO (P-FOXO) and P-ERK. Consistent with western blot analyses, we observed increased P-FOXO only in the DMAQ-B1-treated cells and P-ERK in both individual and combined-treated cells (**Fig. 1E**). Further, P-FOXO localization was primarily cytosolic (**Fig. 1F**) in DMAQ-B1 treated cells and P-ERK localization was primarily nuclear (**Fig. 1G**) in AKT inhibitor and combined treated cells, confirming that the transcription factors involved in RNAi and JAK/STAT induction are both nuclear and transcriptionally active under the expected treatment conditions. We also observed increased transcript expression of *AGO2* and *virus-induced RNA-1* (*vir-*1) which are indicative of RNAi and JAK/STAT activation, respectively, in cells treated with the combined drugs (**Fig. 1H-I**). Collectively, these data indicated that DMAQ-B1 and AKT inhibitor VIII treatment alter IIS phosphorylation in *Ae. aegypti* cells in a pattern consistent with the activation of FOXO- and ERK-dependent antiviral signaling.

**Figure 1:**
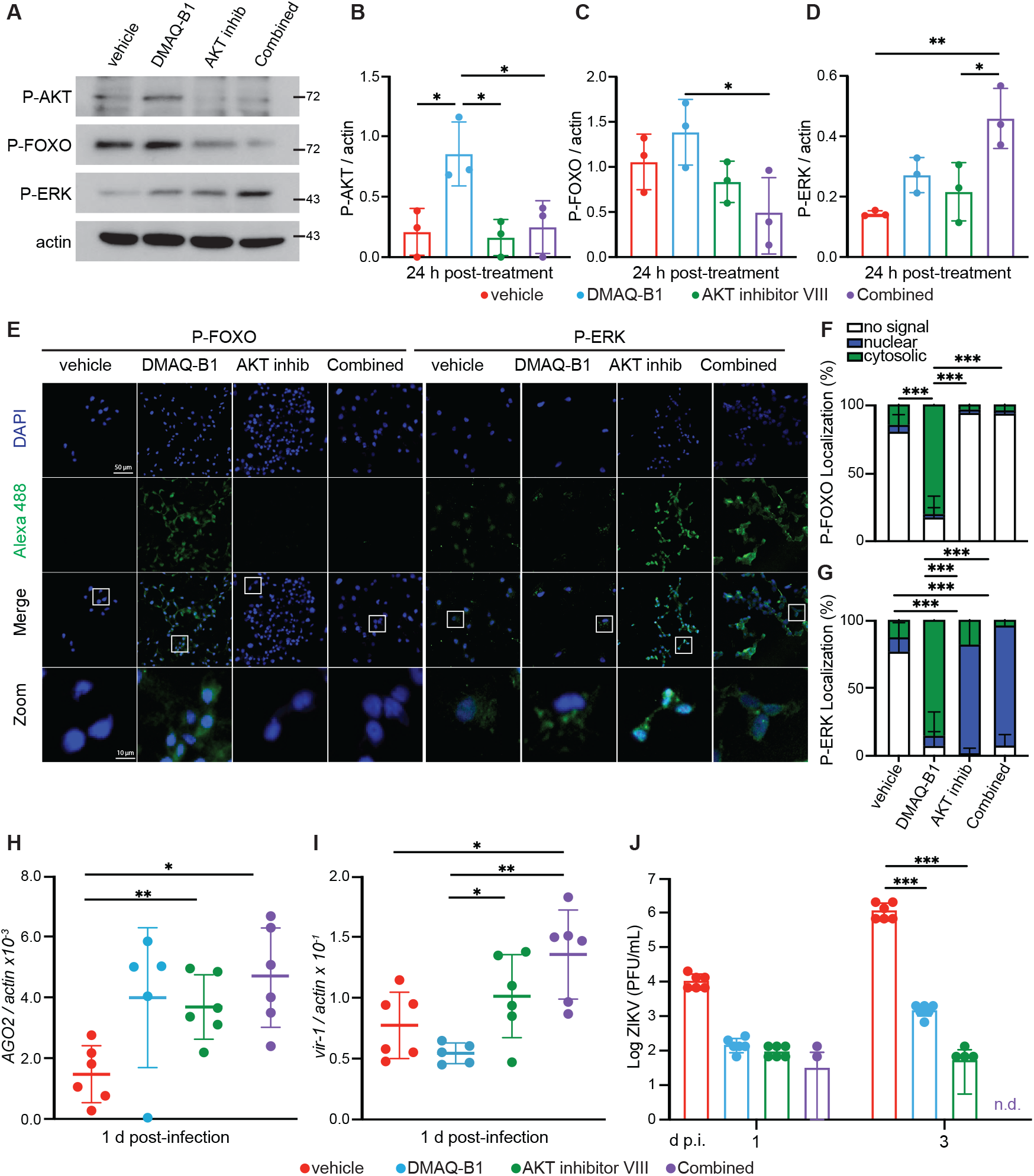
DMAQ-B1 and AKT inhibitor VIII activated RNAi and JAK/STAT *in vitro*. Aag2 cells were treated with vehicle (DMSO), 1 μM DMAQ-B1, 10 μM AKT inhibitor VIII, or combined drugs for 24 h. (A-D) Phosphorylation of AKT, FOXO, and ERK were measured by western blot and phosphorylation was quantified for (B) P-AKT, (C) P-FOXO, and (D) P-ERK by densitometry and normalized to actin (*p < 0.05, One-way ANOVA with Tukey’s test correction for multiple comparisons). (E) P-FOXO and P-ERK abundance and localization were visualized in DAPI-stained Aag2 cells by immunofluorescence microscopy. (F-G) Protein localization of (F) P-FOXO and (G) P-ERK was quantified in microscopy images to evaluate whether fluorescent-tagged proteins was cytosolic or nuclear within individual cells by manually counting cells in images (***p<0.001, Two-way ANOVA with Tukey’s correction). (H-I) Induction of RNAi and JAK/STAT signaling was evaluated as transcript levels of (H) *AGO2* and (I) *vir-1* by qRT-PCR (*p < 0.05; **p<0.01, Unpaired t test with Welch’s correction for multiple comparisons). (J) Aag2 cells that received vehicle, individual, or combined drug treatment for 24 h were infected with ZIKV (MOI 0.01) and supernatant was collected at 1 and 3 dpi. Supernatant virus was titered by standard plaque assay (***p<0.001, Two-way ANOVA with uncorrected Fisher’s LSD). Closed circles represent individual replicates. Horizontal bars represent mean and error bars represent SD. Results are representative of triplicate independent experiments.

Based on these effects on RNAi and JAK/STAT signaling in the absence of virus, we sought to determine the effects of single and combined drugs on ZIKV replication in *Ae. aegypti* cells. Aag2 cells were primed with individual and combined drugs for 24 h prior to infection with the clinically isolated PRVABC59 strain of ZIKV (**Fig. 1J**). We observed significant reductions in ZIKV titer in cells treated with individual and combined drugs by 3 days post-infection (dpi). Most notably, ZIKV titers were undetectable by 3 dpi in cells treated with the combined drugs (**Fig. 1J**). Patel and Hardy (2012) showed that AKT inhibitor VIII was antiviral in Sindbis virus (SINV)-infected *Aedes albopictus* C6/36 cells, but the dysfunctional RNAi response in these cells (Brackney et al., 2010) precluded the confirmation of mechanism in its entirety. Accordingly, we concluded that DMAQ-B1 and AKT inhibitor VIII treatments induced an antiviral response that was increased to the point of non-detectable ZIKV titers when these treatments were combined.

### *DMAQ-B1 and AKT inhibitor VIII treatment of ZIKV-infected* Aedes aegypti *induced simultaneous activation of RNAi and JAK/STAT signaling*

Based on IIS-dependent antiviral activity of DMAQ-B1 and AKT inhibitor VIII *in vitro*, we sought to evaluate whether similar drug effects could be detected in *Ae. aegypti* adult females. Aged-matched 6–9-day old female mosquitoes were fed a ZIKV-containing bloodmeal including vehicle, individual, or combined 10 μM DMAQ-B1 and 10 μM AKT inhibitor VIII. Drug concentrations were selected based on mortality studies to measure drug lethality to mosquitoes over a dose range (**Fig. S2**). Mosquitoes were collected at 3, 7, and 11 dpi, timepoints that corresponded with complete digestion of the blood meal, progression of viremia into distal tissues, and virus infection of the salivary glands (Roundy et al., 2017; Weger-Lucarelli et al., 2016; Williams et al., 2020). Expression levels of RNAi and JAK/STAT signaling gene products were quantified by qRT-PCR at 7 dpi (**Figs. 2A-D**) and 11 dpi (**Fig. 2E-H**). *AGO2* and *p400* were examined as markers of RNAi (**Figs. 2A-B, 2E-F**) (Bernhardt et al., 2012; McFarlane et al., 2020), while *Vago2* and *vir-1* were examined as downstream effectors of JAK/STAT (**Figs. 2C-D, 2G-H**) (Asad et al., 2018; Diop et al., 2019). As in Aag2 cells, we observed that the combination drug treatment resulted in higher expression of RNAi and JAK/STAT signaling gene products at 7dpi (**Figs. 2A-D**). Interestingly, at 11 dpi, only transcript levels for *AGO2* remained significantly higher for individual drug- and combination drug-treated mosquitoes (**Figs. 2E-H**). Based on high transcript expression at 3 dpi in mosquitoes treated with the drug combination (**Fig. S3**), loss of gene induction between 7 and 11 dpi suggests that drug treatment may have a limited efficacy by 11 days-post bloodmeal.

**Figure 2:**
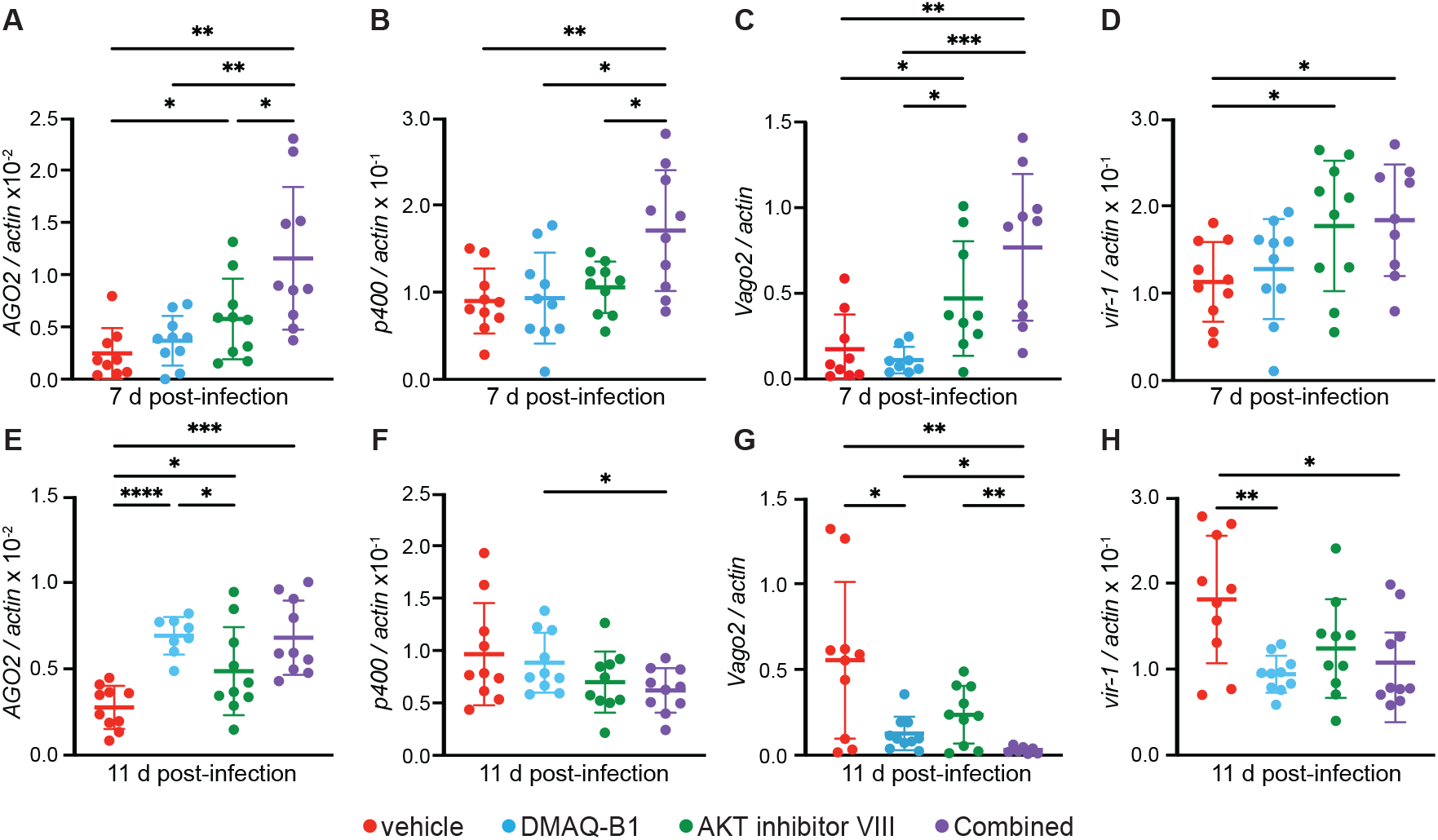
Combined drug treatment induced activation of RNAi and JAK/STAT in *Aedes aegypti* at 7 dpi that was reduced by 11 dpi. Induction of RNAi and JAK/STAT gene transcripts at (A-D) 7 dpi and (E-H) 11 dpi was measured in whole mosquitoes infected with ZIKV and treated with vehicle, 10 μM DMAQ-B1, 10 μM AKT inhibitor VIII, or combined drugs. Transcripts were measured for (A, E) *AGO2*, (B, F) *p400*, (C, G) *Vago2*, and (D, H) *vir-1* by qRT-PCR (*p < 0.05; **p<0.01; ***p<0.001; **** p<0.0001, Unpaired t test with Welch’s correction for multiple comparisons). Closed circles represent individual replicates. Horizontal bars represent mean and error bars represent SD. Results represent duplicate independent experiments.

### *DMAQ-B1 and AKT inhibitor treatment reduced infection prevalence and ZIKV titer in* Aedes aegypti

We next sought to evaluate the effects of individual and combined drug treatments on infection prevalence and ZIKV titers in adult mosquitoes. Mosquitoes were fed a ZIKV-containing bloodmeal treated with vehicle, DMAQ-B1, AKT inhibitor VIII, or combined drug treatment as described. We collected mosquitoes at 3, 7, and 11 dpi and analyzed individual midguts, pairs of salivary glands, and carcasses for ZIKV titers (n=30). There were no differences in virus infection prevalence or viral titers at 3 dpi (**Fig S4**). However, by 7 dpi (**Figs. 3A-C**) and 11 dpi (**Figs. 3G-I**), infection prevalence was reduced relative to vehicle control in mosquitoes treated with individual and combined drug treated mosquitoes. Viral titers in ZIKV-positive mosquitoes were reduced relative to vehicle control less than two-fold at 7 dpi (**Figs. 3D-F**), but this reduction was greater than two-fold at 11 dpi (**Figs. 3J-L**). Notably, infection prevalence and salivary gland viral titers were reduced in combined drug-treated mosquitoes at 11 dpi, a time consistent with virus transmission during feeding (Armstrong et al., 2020; Sánchez-Vargas et al., 2018).

**Figure 3:**
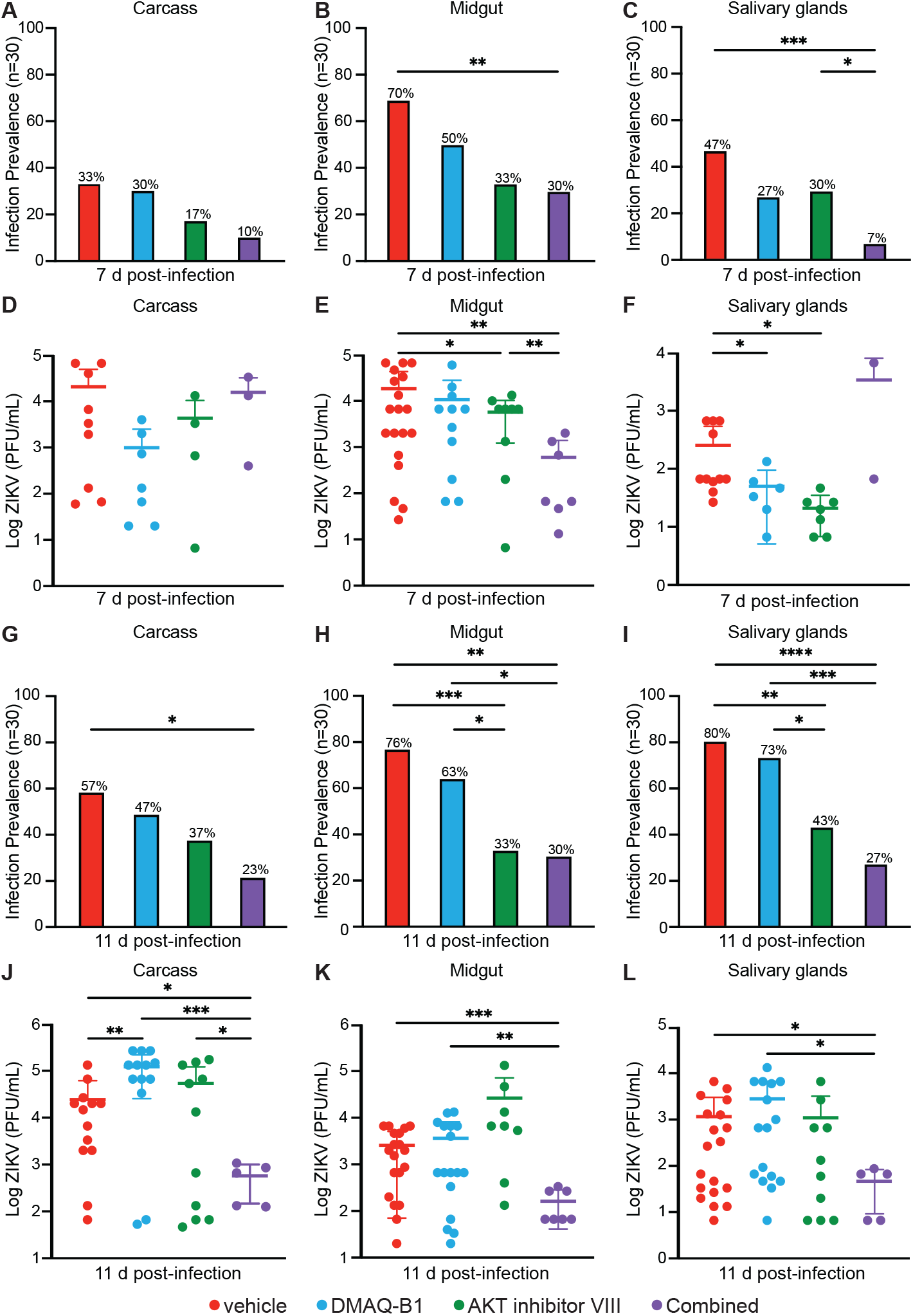
Individual and combined drug treatments reduced infection prevalence and ZIKV titers in *Aedes aegypti*. Individual mosquito midguts, pairs of salivary glands, and carcasses (n=30) were titered for ZIKV by standard plaque assay at (A-F) 7dpi and (G-L) 11 dpi. (A-C, G-I) Infection prevalence was calculated as the ratio of ZIKV-positive samples to the total sample size (*p < 0.05; **p<0.01; ***p<0.001, Two-tailed Fisher’s exact test). (D-F, J-L) Viral titers were determined in ZIKV-positive mosquitoes (*p<0.05; **p<0.01; ***p<0.001; **** p<0.0001, Unpaired t test with Welch’s correction for multiple comparisons).

Mosquitoes that received combined drug treatment were not only less likely to be ZIKV-positive, but among those individuals that were ZIKA-positive, salivary gland viral load was substantially reduced. Average gland viral load at 11 dpi (1.7log10±1.5 PFU/mL) was below titers previously associated with successful virus transmission (4.8log10±0.6 PFU/mL) (Vazeille et al., 2019). These observations suggest that combined drug treatment and coordinated activation of RNAi and JAK/STAT provides antiviral immunity against ZIKV that effectively reduced infection prevalence and viral titers below reported transmissible levels. These effects of combined drug treatment would be predicted, therefore, to reduce mosquito transmission of ZIKV.

### Inhibition of RNAi and JAK/STAT signaling resulted in loss of drug-mediated antiviral protection

To confirm that DMAQ-B1 and AKT inhibitor VIII mediate RNAi- and JAK/STAT-dependent antiviral responses, we transfected Aag2 cells with siRNA constructs to knockdown expression of *AGO2* and *vir-1*. Cells were transfected with siRNAs that targeted either gene individually (siAGO2, siVir-1) or stacked gene expression (siAGO2 + siVir-1) (Terradas et al., 2017). We observed significantly reduced gene expression at 48 h post transfection for both individual and stacked siRNA treatments compared to cells that were treated with control, non-targeting siRNAs (Clemons et al., 2011) Individual siRNA treated cells exhibited a 69% and 68% reduction in *vir-1* and *AGO2*, respectively, compared to the control siRNA-treated cells. Stacked siRNA treated cells exhibited a 79% and 84% reduction in *vir-1* and *AGO2*, respectively, compared to control siRNA-treated cells (**Figs. 4A and 4B**). Next, we treated cells at 48 h after siRNA transfection with vehicle, individual, or combined drug treatments for 24 h prior to ZIKV infection. Viral titers were measured in supernatants collected at 2 dpi to determine if antiviral protection was impacted in the absence of antiviral RNAi, JAK/STAT, or both. We observed that both individual and combined *AGO2* and *vir-1* knockdowns resulted in significant losses of drug-mediated antiviral protection relative to drug-treated controls (**Fig. 4C**). Interestingly, while we observed a loss in antiviral protection between pair-wise treatment conditions, we did not observe a significant difference in viral titers among siRNA transfections. These results suggested that IIS-dependent RNAi and JAK/STAT signaling are sufficient to significantly reduce ZIKV titers, but other pathways may also contribute to this biology. For example, despite the reduction in ZIKV to undetectable levels via IIS-dependent antiviral immunity, Toll signaling (Angleró-Rodríguez et al., 2017) and autophagy (Liu et al., 2018) could contribute to control of ZIKV replication. Collectively, our data suggest that repurposing small molecule drugs to target mosquito IIS can induce antiviral responses that significantly reduce ZIKV infection prevalence and transmission potential in *Ae. aegypti* through activation of RNAi and JAK/STAT signaling.

**Figure 4:**
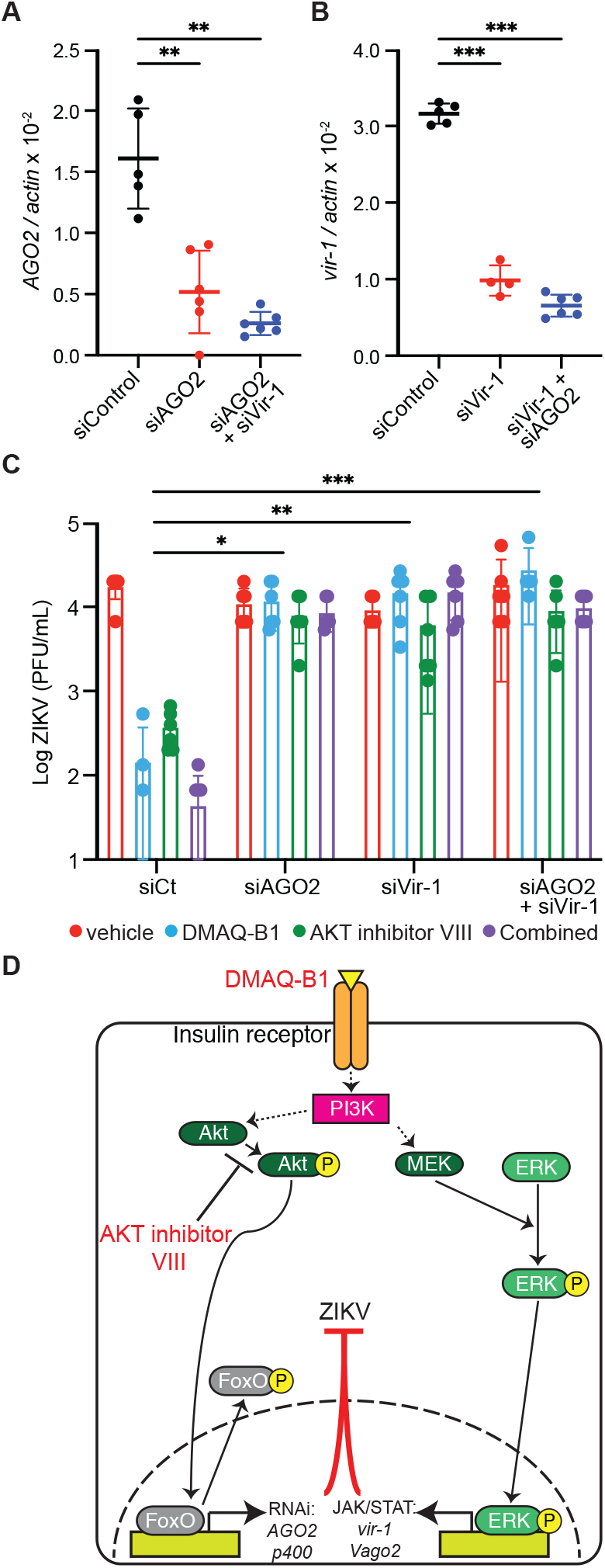
Knockdown of RNAi and JAK/STAT signaling resulted in loss of drug-mediated antiviral protection. (A-B) AGO2 and vir-1 were knocked down in Aag2 cells and transcript levels were determined for (A) *AGO2* and (B) *vir-1* by qRT-PCR for cells transfected with scramble control (siControl), individual siRNA construct, or stacked siRNA (siAGO2+siVir-1) (**p<0.01; ***p<0.001; **** p<0.0001, Unpaired t test with Welch’s correction). (C) 48 h following transfection, cells were primed with DMAQ-B1 or AKT inhibitor VIII for 24 h prior to infection with ZIKV (MOI=0.01 PFU/cell). Supernatant was collected at 2 dpi and virus was titered by standard plaque assay (*p<0.05; **p<0.01; ***p<0.001 Two-way ANOVA with uncorrected Fisher’s LSD test). (D) Schematic of proposed mechanism of action mediated by DMAQ-B1 and AKT inhibitor VIII during ZIKV infection through simultaneous induction of antiviral RNAi and JAK/STAT signaling.

## DISCUSSION

Global climate change has enabled the expansion of mosquito populations into new ranges with concomitant increases in the variety and incidence of mosquito-borne diseases. Recent ZIKV epidemics have demonstrated the need for host-virus interactions research to identify novel drug targets and for the development of more effective means of vector control. Current vector control efforts involving microbiota or genetic manipulations, while promising, could be enhanced by the addition of antiviral drug strategies to ongoing control efforts.

In the present work, we evaluated the therapeutic potential of IIS-targeted small molecules to reduce ZIKV infection prevalence and titers in *Ae. aegypti*. We demonstrated that the potent insulin mimetic DMAQ-B1 and the AKT inhibitor VIII synergized IIS-mediated antiviral immunity in *Ae. aegypti* to reduce ZIKV infection prevalence and titers in infected mosquitoes (**Fig. 4D**). While this study is not the first to identify IIS regulation of antiviral immunity, we have advanced this field by demonstrating that readily available and potent IIS-targeted small molecules induced substantial and significant antiviral immunity in *Ae. aegypti* against a clinically virulent strain of ZIKV. By targeting IIS as a mediator of two independent antiviral pathways, we reduced not only infection prevalence but also virus titers below levels previously associated with successful transmission. Both of these effects would be predicted to reduce ZIKV transmission by the primary vector *Ae. aegypti*. In demonstrating these effects, we have also provided a foundation for future translation of our findings to the field. Specifically, we seek to advance small molecule delivery via attractive bait stations to induce IIS-mediated, broad antiviral immunity in mosquitoes that ingest these compounds.

Of particular interest for a potential field-based strategy is the broad impact that IIS appears to have across species. Previous studies have confirmed that exogenous treatment with or the endogenous effects of insulin in *D. melanogaster* and *Culex* spp. reduced replication of both WNV and DENV (Ahlers et al., 2019; Xu et al., 2013). Interestingly, the antiviral effects of IIS during ZIKV infection are not limited to arthropod species. In mammalian models, ZIKV NS4A/NS4B activates PI3K-AKT signaling that is associated with neurogenetic dysregulation (Liang et al., 2016). Further, the broadly antiviral celecoxib kinase inhibitor AR-12 and AKT inhibitor VIII have been shown to reduce ZIKV replication and pathogenesis in mice by blocking PI3K-AKT activation (Chan et al., 2018). It is also established that diabetic individuals with dysfunctional IIS are more susceptible to severe disease during WNV (Kumar et al., 2012, 2014), DENV (Lee et al., 2020), and ZIKV infection (Nielsen and Bygbjerg, 2016). Given the variety of mosquito species that deploy IIS-dependent immunity against notable major arboviruses, it would be worth investigating whether similarly broad IIS regulation of antiviral responses can be detected in mammalian hosts. If so, it may be possible to develop IIS-targeted transmission blocking therapeutic drugs that mitigate Zika disease and, when delivered in blood from treated patients to *Ae. aegypti*, reduce infection in and transmission by the mosquito vector.

## ACKNOWLEDGEMENTS

We thank A. Nicola, S. O’Neill, G. Ebel, and K. Olson for cells, viruses, and mosquitoes used in these experiments. We thank G. Terradas and E. McGraw for providing siRNA sequences. We also like to thank S. Bennett, C. Blair, and B. Foy, amongst others of the Center for Vector-borne Infectious Diseases, for assistance with mosquito husbandry and feedback regarding experimental design. We also thank L.R.H Ahlers for critical reading of our manuscript. This research was supported by the WSU College of Veterinary Medicine Stanley L. Adler research fund to A.G.G, NIH / National Institute of General Medical Sciences (NIGMS)-funded pre-doctoral fellowship (T32 GM008336) and a Poncin Fellowship to C.E.T., NIH/NIAID grant R01AI151166 to I.S.V. and R.P., and University of Idaho and UI College of Agricultural and Life Sciences startup funds to S.L..

## AUTHOR CONTRIBUTIONS

Conceptualization, C.E.T. and A.G.G.; Methodology, C.E.T., G.R., I.S.V., R.P., S.L., and A.G.G.; Validation, C.E.T., G.R., and I.S.V.; Investigation, C.E.T., G.R., I.S.V., R.P., and A.G.G; Resources, I.S.V., R.P., S.L., and A.G.G.; Writing – Original Draft, C.E.T.; Writing – Review and Editing, G.R., I.S.V., R.P., S.L., and A.G.G.; Visualization, C.E.T. and A.G.G.; Funding Acquisition, C.E.T., I.S.V., R.P., S.L., and A.G.G.

## DECLARATION OF INTERESTS

The authors declare no competing interests.

## MATERIALS AND METHODS

### CONTACT FOR REAGENT AND RESOURCE SHARING

Further information and requests for resources and reagents should be directed to and will be fulfilled by the Lead Contact, Alan Goodman.

### EXPERIMENTAL MODEL AND SUBJECT DETAILS

#### Mosquito rearing

*Aedes aegypti* strain Poza Rica, from the state of Veracruz, Mexico were originally collected in 2012, and maintained as described (Weger-Lucarelli et al., 2016). Adult mosquitoes were provided continuous access to water and 10% sucrose *ad libitum*, and the females were allowed to feed on defibrinated sheep blood (Colorado Serum Company) supplemented with 1mM ATP using an artificial feeding system to stimulate oogenesis.. Larvae were reared and maintained under constant 28 °C, 70% humidity, and 12-hour light, 12 hour dark diurnal cycle. 6-9 day old adult female mosquitoes were deprived of sucrose 24 hours prior to experimental feedings as described (Weger-Lucarelli et al., 2016). Mosquito infections, maintenance, and plaque assays were performed under BSL3 and ACL3 facilities, approved by Colorado State University’s Institutional Biosafety Committee 16-074B.

#### Cells and virus

Vero cells (ATCC, CRL-81) were provided by A. Nicola and cultured at 37 °C/5% CO_2_ in DMEM (ThermoFisher 11965) supplemented with 10% FBS (Atlas BiologicalsFS-0500-A) and 1x antibiotic-antimycotic (ThermoFisher 15240062). *Ae. aegypti* Aag2 cells (*Wolbachia*-free) (Terradas et al., 2017) were gifted by S. O’Neill and cultured as described (Terradas et al., 2017). For drug treatment, culture media with 2% FBS were supplemented with 1% DMSO, 1 μM DMAQ-B1, 10 μM AKT inhibitor VIII, or combined drugs.

Concentrations of DMAQ-B1 and AKT inhibitor VIII were selected at non-cytotoxic levels for both cell culture (**Figure S1**) and adult mosquitoes (**Figure S2**). ZIKV strain PRVABC59 (Accession # KU501215) was obtained from the CDC and was isolated in 2015 from a clinical case in Puerto Rico and prepared as described (Weger-Lucarelli et al., 2016).

### METHOD DETAILS

#### *In vitro* replication

Aag2 cells were seeded into a 24-well plate at a confluency of 5×10^5^ cells/well with 6 independent wells for each experimental condition. The following day, cells were treated with 1% DMSO, 1 μM DMAQ-B1, 10 μM AKT inhibitor VIII, or combined drugs in 2% FBS media as described (Ahlers et al., 2019) for 24 h prior to infection. Cells were then infected with ZIKV at MOI of 0.01 PFU/cell for 1 h. Virus inoculum was removed, and fresh experimental media was added. Supernatant samples were collected at 1 and 3 dpi for later titration. ZIKV titers were determined by standard plaque assay on Vero cells (Baer and Kehn-Hall, 2014; Sanchez-Vargas et al., 2021).

#### Cytotoxicity of DMAQ-B1 and AKT inhibitor VIII

Cytotoxicity of DMAQ-B1 and AKT inhibitor VIII was evaluated in both cell culture and in adult female *Ae. aegypti*. DMAQ-B1 and AKT inhibitor VIII was added to a monolayer of 2.5×10^5^ cells/well in 48-well plates at various concentrations (100 μM, 10 μM, 1 μM, 0.1 μM). Cells were collected at 1, 2, and 3 d post-treatment, stained with trypan blue (ThermoFisher 15250-061) and scored as live or dead as described (Ahlers et al., 2016). Combined DMAQ-B1 and AKT inhibitor VIII cytotoxicity was evaluated using the maximum individual concentrations that corresponded to minimal cytotoxicity. A total of eight technical replicates were averaged for each biological replicate. 1% DMSO treated and 1% Triton X-100 treated cells were also scored as negative and positive controls, respectively. Cytotoxicity was evaluated similarly in 6-9 day old female mosquitoes exposed to a bloodmeal containing small molecule drugs (100 μM, 10 μM, 1 μM), 1% DMSO vehicle control, or blood only. Following 1 h of feeding, engorged females were kept and maintained on sucrose for 14 d to monitor mortality. Combined small molecule drug treatment was evaluated using observed lethal and nonlethal individual concentrations. Each experimental group contained approximately 70-100 mosquitoes.

#### Immunoblotting

Protein extracts were prepared by lysing cells with RIPA buffer (25 mM Tris-HCl pH 7.6, 150 mM NaCl, 1 mM EDTA, 1% NP-40, 1% sodium deoxycholate, 0.1% SDS, 1mM Na_3_VO_4_, 1 mM NaF, 0.1 mM PMSF, 10 μM aprotinin, 5 μg/mL leupeptin, 1 μg/mL pepstatin A). Protein samples were diluted using 2x Laemmli loading buffer, mixed, and boiled for 5 minutes at 95 °C. Samples were analyzed by SDS/PAGE using a 10% acrylamide gel, followed by transfer onto PVDF membranes (Millipore IPVH00010). Membranes were blocked with 5% BSA (ThermoFisher BP9706) in Tris-buffered saline (50 mM Tris-HCl pH 7.5, 150 mM NaCl) and 0.1% Tween-20 for 1 h at room temperature.

Primary antibody labeling was completed with anti-P-Akt (1:1,000; Cell Signaling 4060), anti-P-ERK (1:1000; Sigma M8159), anti-P-FOXO (1:1000; Millipore 07-695), or anti-actin (1:10,000; Sigma A2066) overnight at 4 °C. Secondary antibody labeling was completed using anti-rabbit IgG-HRP conjugate (1:10,000; Promega W401B) or anti-mouse IgG-HRP conjugate (1:10,000; Promega W402B) by incubating membranes for 2 h at room temperature. Blots were imaged onto film using luminol enhancer (ThermoFisher 1862124). Densitometry analysis was completed using three independent blots using BioRad Image Lab with bands normalized to actin.

#### RNA interference *in vitro*

Long dsRNA targeting *Ae. aegypti AGO2, vir-1*, and non-targeting control dsRNA was synthesized as described (Terradas et al., 2017). Targeted sequences and primers are listed in **Table S1**. dsRNA was transfected into Aag2 cells as described (Terradas et al., 2017) for 48 h prior to small molecule treatment and infection. RNA was extracted and purified to confirm reduced expression by qRT-PCR and viral concentration was confirmed by standard plaque assay.

#### Quantitative reverse transcriptase PCR

qRT-PCR was used to measure mRNA levels in *Ae. aegypti* Aag2 cells and adult females. Cells or mosquitoes were lysed with Trizol Reagent (ThermoFisher 15596). RNA was isolated by column purification (ZymoResearch R2050), DNA was removed (ThermoFisher 18068), and cDNA was prepared (BioRad 170–8891). Expression of *Ae. aegypti AGO2, p400, Vago2*, and *vir-1* were measured using SYBR Green reagents (ThermoFisher K0222) and normalized to *actin*. The reaction for samples included one cycle of denaturation at 95 °C for 10 minutes, followed by 45 cycles of denaturation at 95 °C for 15 seconds and extension at 60 °C for 1 minute, using an Applied Biosystems 7500 Fast Real Time PCR System. ROX was used as an internal control. qRT-PCR primer sequences are listed in **Table S1**.

#### Immunofluorescence microscopy

*Ae. aegypti* Aag2 cells were seeded onto coverslips in 12-well plates at a confluency of approximately 1×10^6^ cells/well. Cells were then treated for 24 hours with 1% DMSO, 1 μM DMAQ-B1, 10 μM AKT inhibitor VIII, or combined small molecule treatment supplemented in 2% FBS media as described (Ahlers et al., 2019). Coverslips were fixed in 4% paraformaldehyde for 10 minutes at room temperature, permeabilized in 0.1% Triton-X-100 for 30 minutes at room temperature and blocked in 1% BSA in TBS for 30 min at 37 °C. Primary antibody labeling was completed with anti-P-FOXO (1:100) and anti-P-ERK (1:100) for 2 h at humified room temperature. Secondary antibody labeling was completed using anti-rabbit (Life Technologies A11034) or anti-mouse (Life Technologies A11029) Alexafluor 488 (1:300) by incubating membranes for 1 h at room temperature in the dark. Samples were stained with DAPI (1:100; Cell Signaling 4083), mounted onto coverslips using ProLong Diamond Antifade Mountant (Invitrogen P36961), and imaged using a Leica Sp8X confocal microscope. Localization percentages were determined by counting the total number of cells and evaluating if green-fluorescent signal was cytosolic, nuclear, or no signal in relation to DAPI-stained nuclei.

#### Mosquito infections

Fresh ZIKV virus stock was made from Vero cells infected at MOI of 0.1 PFU/cell at 72 hours prior to bloodmeal feed of 6-9 day old female mosquitoes as described (Weger-Lucarelli et al., 2016; Williams et al., 2020). Mosquitoes were fed a bloodmeal supplemented with 1 mM ATP and infected with the fresh virus inoculum that was back-titrated to 2.4×10^6^ PFU/mL. Bloodmeals included 1% DMSO vehicle control, 10 μM DMAQ-B1, 10 μM AKT inhibitor VIII, or combined drugs. Following 1 h of feeding, mosquitoes were anesthetized on ice and engorged mosquitoes were moved into new cartons and maintained on sucrose. Mosquitoes were collected at 3, 7, and 11 dpi in which the midgut, salivary glands, and carcass were separated, homogenized, and filtered for infection determination and viral titering. Whole mosquitoes were collected at the same timepoints for qRT-PCR analysis.

## QUANTIFICATION AND STATISTICAL ANALYSIS

Results presented as dot plots show data from individual biological replicates (n=3-10) and the arithmetic mean of the data, shown as a black horizontal line. Biological replicates of adult mosquitoes (n=2-24) consisted of two pooled mosquitoes. Results shown are representative of at least duplicate independent experiments, as indicated in the figure legends. All statistical analyses of biological replicates were completed using GraphPad Prism 9 and significance was defined as p<0.05. One-way ANOVA with Tukey’s correction for multiple comparisons was used for densitometry analysis. Two-way ANOVA with Tukey’s correction for multiple comparisons was used for microscopy localization analysis. Unpaired t test with Welch’s correction was used for qRT-PCR and *in vivo* viral titer analysis. Two-way ANOVA with uncorrected Fisher’s LSD test was used for analysis of multiday *in vitro* viral titer. Two-tailed Fisher’s exact test was used to compare infection prevalences. Two-way ANOVA with Tukey’s correction for multiple comparisons was used for analysis of small molecule cytotoxicity *in vitro* and *in vivo*. All error bars represent standard deviation (SD) of the mean. Outliers were identified using a ROUT test (Q=5%) and removed.

## SUPPLEMENTAL INFORMATION

**Table S1.**
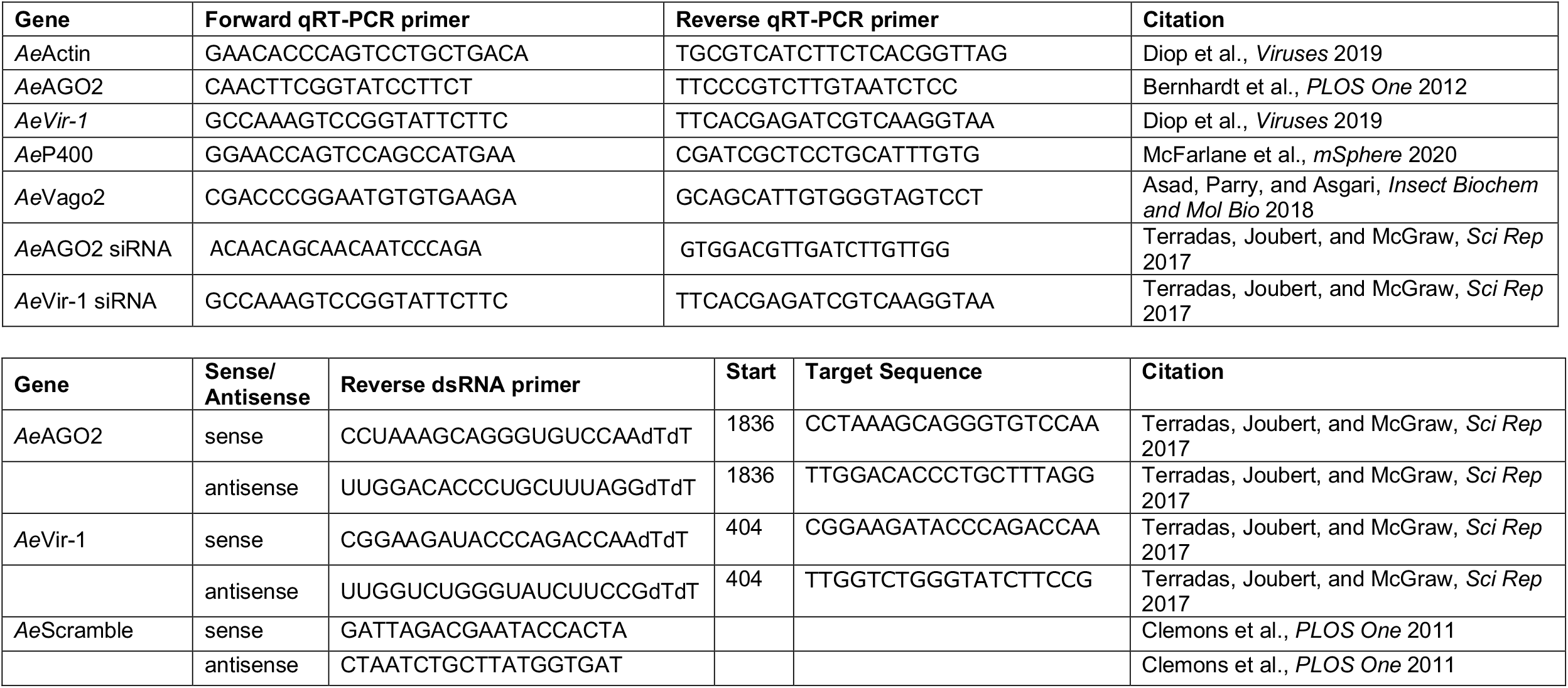
Primers for qRT-PCR and siRNA synthesis.

**Figure S1:**
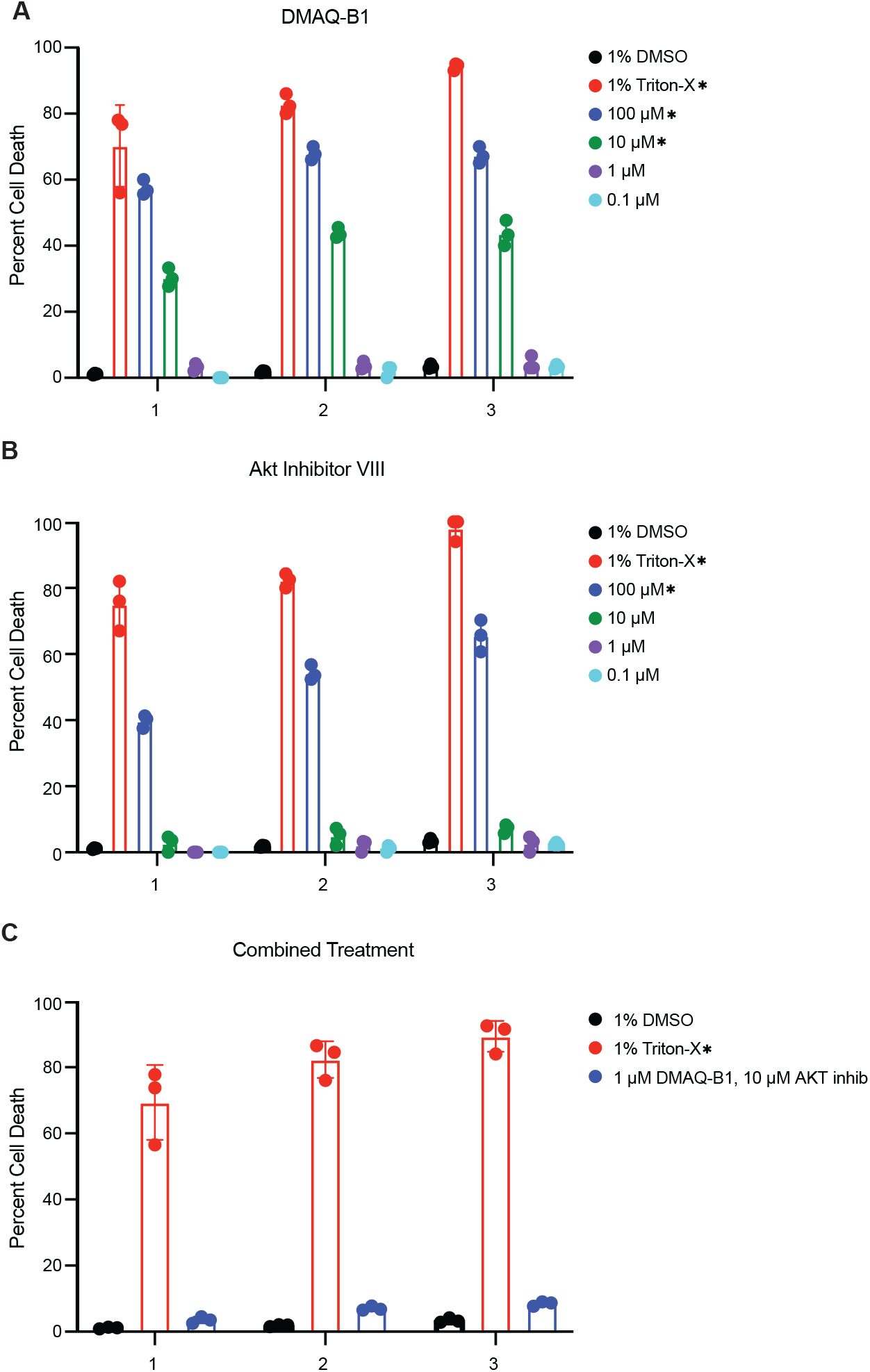
DMAQ-B1 and AKT inhibitor VIII exhibited dose-dependent cytotoxicity in Aag2 cells. Aag2 cells were treated with various concentrations of (A) DMAQ-B1, (B) AKT inhibitor VIII, (C) combined drugs, or DMSO vehicle control and cell viability was measured by trypan blue exclusion. Cells that received 1% Trixton-X-100 treatment were used as a positive, 100% lethality control. Closed circles represent biological replicates measured in technical triplicate. Horizontal black bars represent the mean. Error bars represent SD. Significance was measured by Two-Way ANOVA with 1% DMSO vehicle control (*p<0.01). Data are representative of triplicate independent experiments.

**Figure S2:**
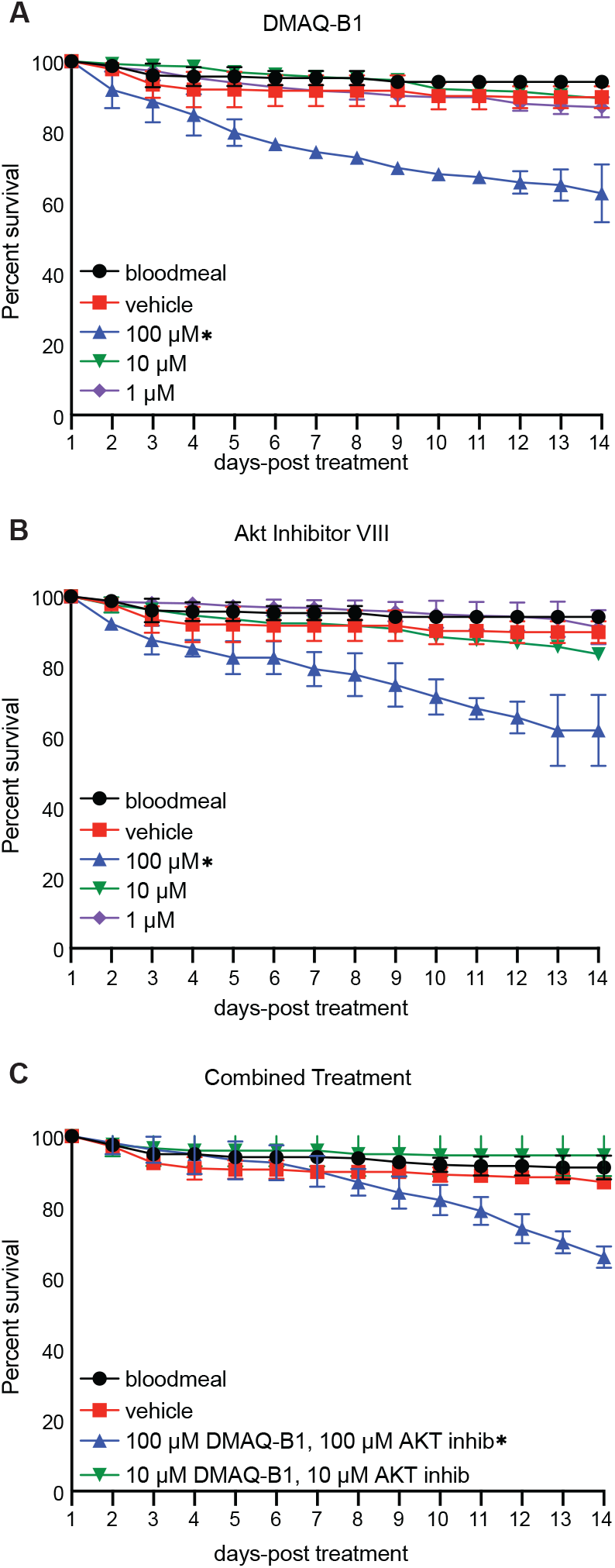
DMAQ-B1 and AKT inhibitor VIII exhibited minimal, dose-dependent toxicity to *Ae. aegypti*. Adult female *Ae. aegypti* were treated with various concentrations of (A) DMAQ-B1, (B) AKT inhibitor VIII, (C) combined drugs and toxicity was measured by survival over 14 days. Closed circles represent percent survival of mosquitos (n=60-100) measured in triplicate. Horizontal black bars represent the mean. Error bars represent SD. Significance was measured by Two-Way ANOVA with 1% DMSO vehicle control (*p<0.05). Data are representative of duplicate independent experiments.

**Figure S3:**
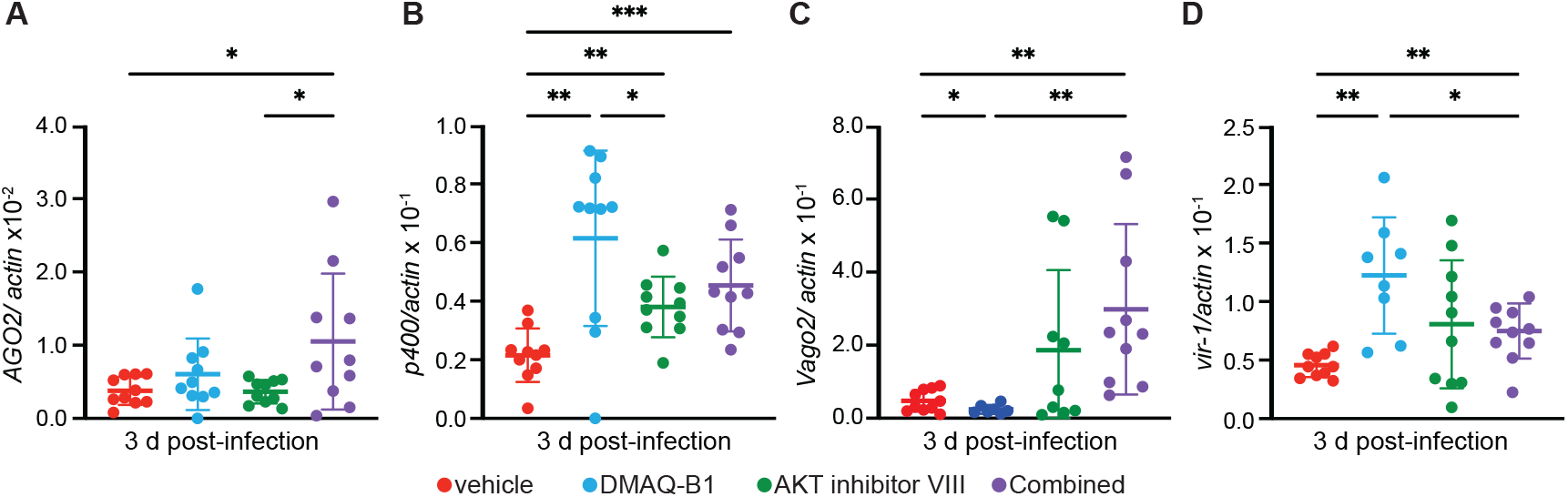
RNAi and JAK/STAT signaling was induced in small molecule treated mosquitoes at 3 dpi. Induction of (A) *AGO2*, (B) *p400*, (C) *Vago2*, and (D) *vir-1* in adult female *Ae. aegypti* was measured by qRT-PCR 3 dpi of ZIKV- and drug-containing bloodmeal. (*p<0.05; **p < 0.01; ***p < 0.001). Open circles represent individual biological replicates. Horizontal black bars represent the mean. Error bars represent SDs. Data are representative of duplicate independent experiments.

**Figure S4:**
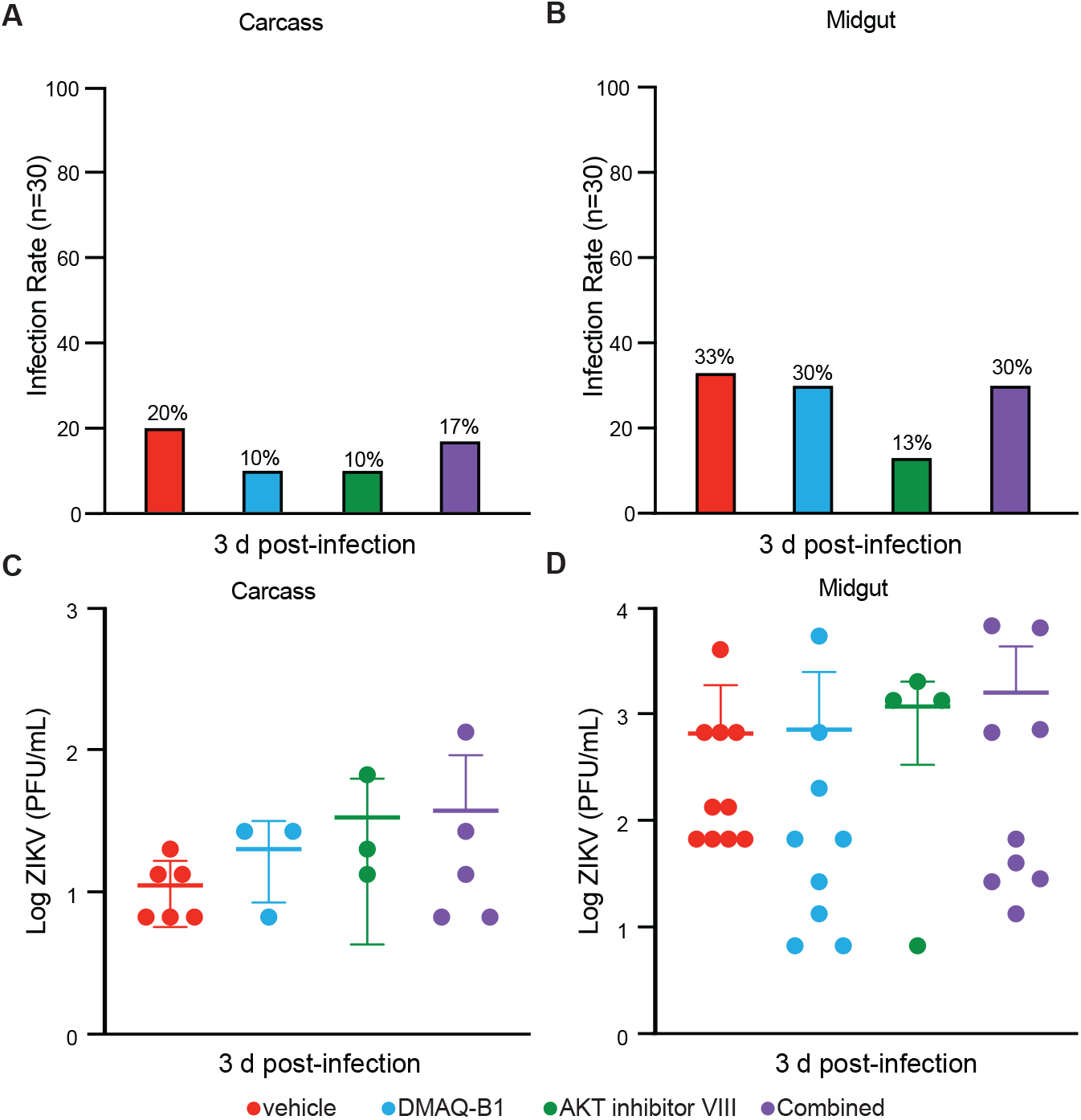
Infection prevalence and ZIKV titers at 3 dpi were not different among small molecule-treated and control *Ae aegypti* at 3 dpi. *Ae. aegypti* were primed with 1% DMSO, 10 μM DMAQ-B1, 10 μM AKT inhibitor VIII, or combined drugs and infected with ZIKV by bloodmeal. Mosquitoes (n=30) were collected at 3 dpi and individual midguts, pairs of salivary glands, and carcasses were prepared and titered by standard plaque assay. Infection prevalence was determined by comparing the number of mosquitoes with detectable virus to the total mosquitoes in the sample. Viral titer was measured in mosquitoes that were positive for ZIKV. There were no differences in infection prevalence or viral titers among conditions. Open circles represent biological replicates. Bars represent the mean. Error bars represent SDs. Data are representative of duplicate independent experiments.

## KEY RESOURCES TABLE

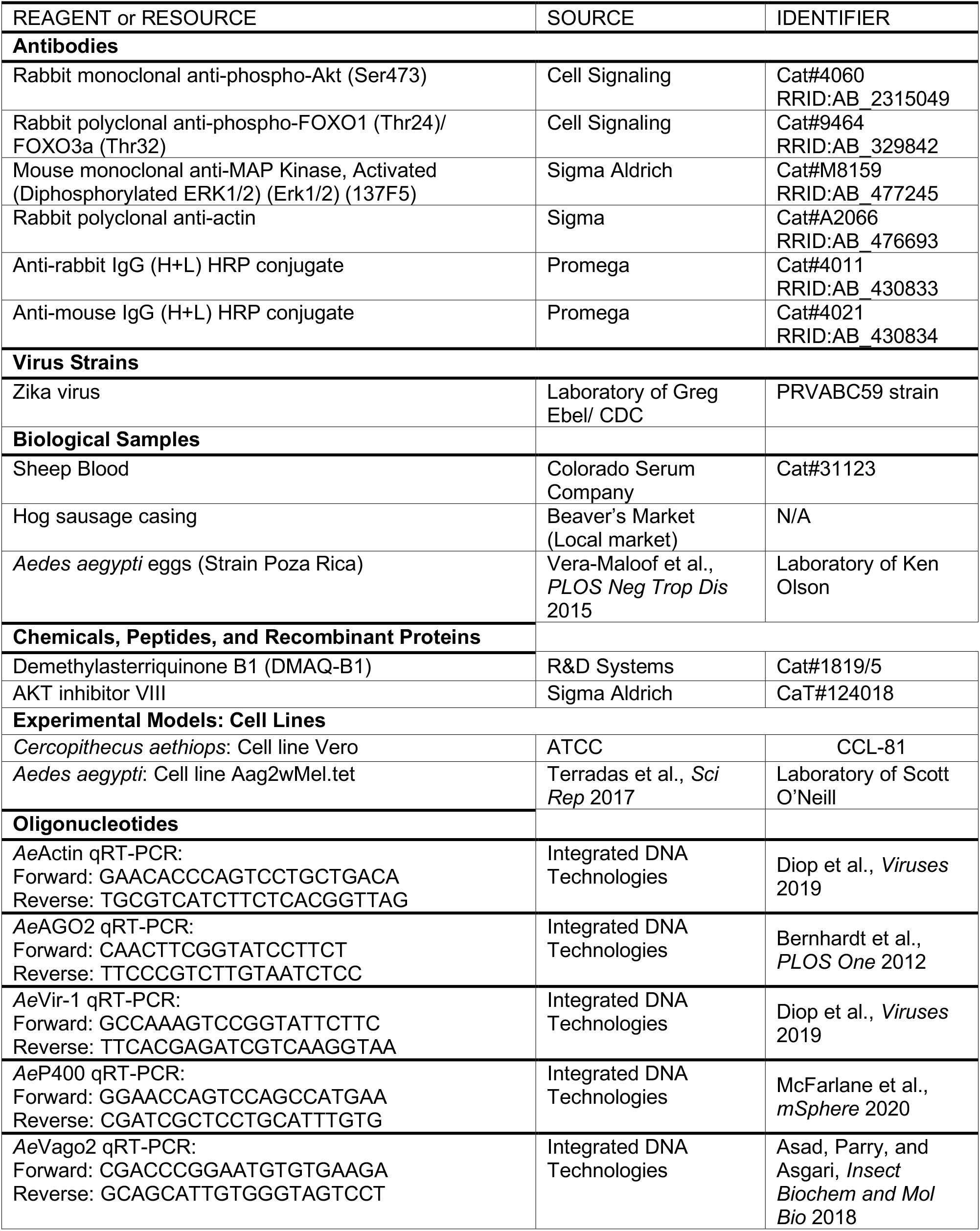
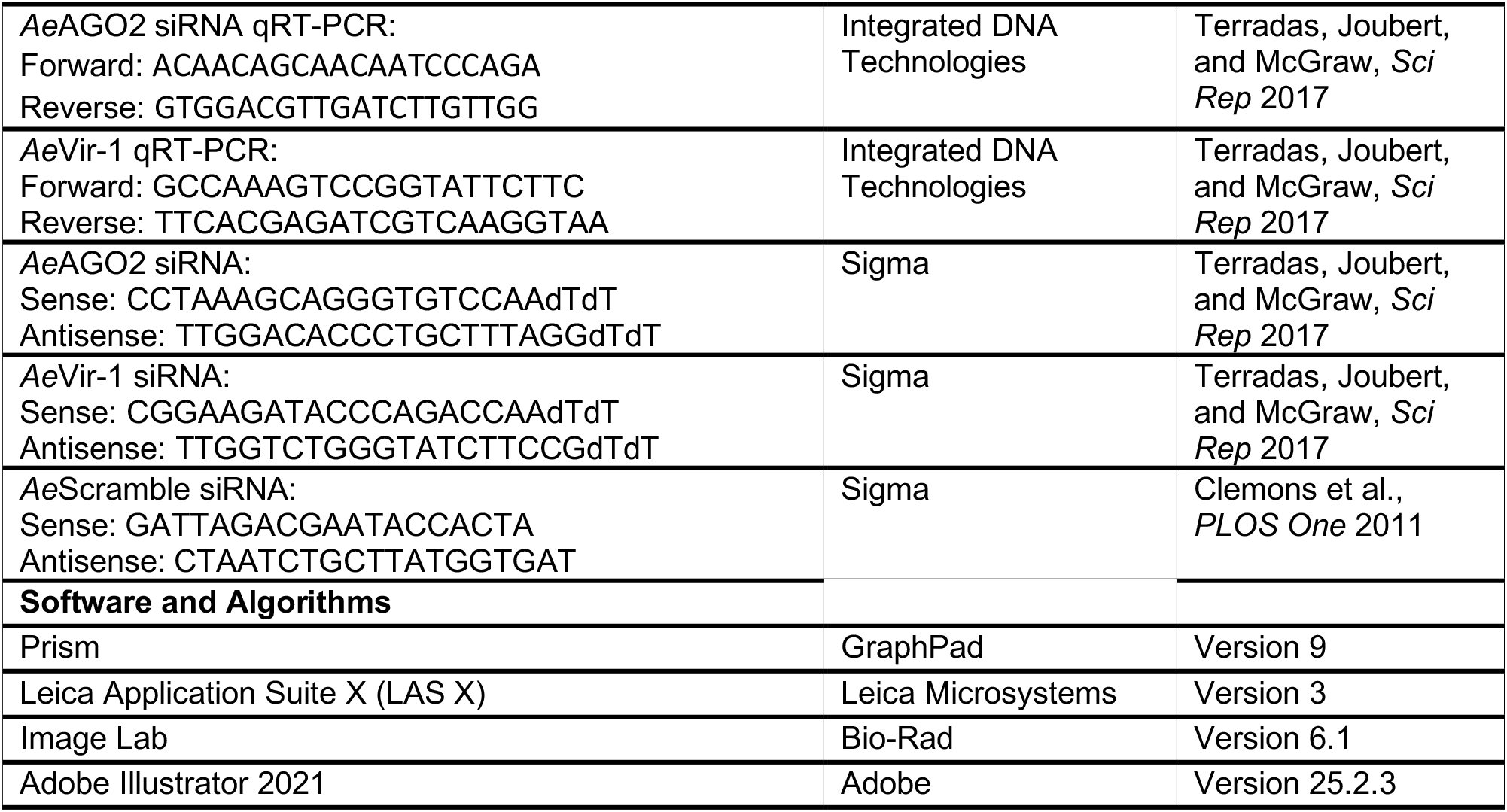

## Notes

### Competing Interest Statement

The authors have declared no competing interest.

